# Adaptive respiratory bypass restores oxidative metabolism in NADH dehydrogenase-deficient *Escherichia coli*

**DOI:** 10.64898/2026.10.01.755914

**Authors:** Arpita Biswas, Arjun Patel, Nikita Goel, Snehal V. Khairnar, Amitesh Anand

## Abstract

Nicotinamide adenine dinucleotide is central to the cellular redox network and continuously cycles between its oxidized (NAD^⁺^) and reduced (NADH) forms. Membrane-bound NADH dehydrogenases are the primary NADH-oxidizing enzymes and principal entry point for reducing equivalents into the bacterial respiratory chain. While maintaining optimal balance of this redox cofactor is among the most fundamental biochemical constraints on living systems, how cells adapt when the primary NADH oxidation route is lost remains poorly understood. Here we show that *Escherichia coli* lacking both NADH dehydrogenases initially compensates through costly fermentative lactate secretion, potentially to maintain redox and energy homeostasis, but at the cost of impaired growth and oxidative metabolism. Using adaptive laboratory evolution across four independent lineages, we found that all evolved populations derepressed malate:quinone oxidoreductase by disrupting an ArcA-binding site. This gain-of-function mutation establishes an alternative pathway that funnels electrons directly into the quinone pool, restoring respiratory oxygen consumption to wild-type level. Multi-omics and genome-scale metabolic modeling reveal that this bypass drives broad reorganization of central carbon metabolism, reinstates tricarboxylic acid cycle flux, and shifts cells toward a higher, more oxidative aero-type. These findings uncover a previously unrecognized metabolic buffering capacity that enables bacteria to circumvent loss of canonical respiratory entry points.

## Introduction

Nicotinamide adenine dinucleotide (NAD) is a crucial redox cofactor, and its evolution traces back to the metabolism of the Last Universal Common Ancestor (LUCA) (1, 2). At least one tenth of all metabolic reactions of *Escherichia coli* directly involve NAD and its phosphorylated analog (NADP), while many additional reactions are indirectly constrained by the cellular NAD(P)^+^/NAD(P)H redox state (Supplementary Sheet 1) (3). In aerobic bacteria, membrane-bound NADH dehydrogenases are the major site for NADH oxidation through which electrons enter the respiratory electron transport system (ETS). These enzymes regenerate NAD^⁺^ and transfer electrons to membrane-bound quinones, coupling redox balance to proton-motive force (PMF) generation (4). To optimize respiratory efficiency under fluctuating environmental conditions, bacteria have evolved distinct classes of NADH dehydrogenases, including the proton-translocating type I NADH dehydrogenase (NDH-I), the non-proton-translocating type II NADH dehydrogenase (NDH-II), and, in some organisms, the sodium-translocating Na^⁺^-NADH:quinone oxidoreductase (NQR) (4, 5). Although these enzymes differ in their energetic efficiency, regulation, and ecological distribution, they collectively perform an indispensable biochemical function by maintaining redox homeostasis and initiating respiratory electron flow.

The central importance of NADH dehydrogenases is reflected in the profound physiological consequences of their disruption. Genetic deletion or pharmacological inhibition of these enzymes elevates intracellular NADH levels, restricts oxidative metabolism, perturbs respiratory energy conservation, and compromises bacterial growth (6–10). In many pathogens, NADH dehydrogenases also influence virulence, persistence, and antimicrobial susceptibility, making them attractive therapeutic targets (11–18). The physiological effects of impaired NADH oxidation have been extensively investigated, yet these studies have largely focused on the consequences of disrupting redox balance at a proximal level, undermining bacterial adaptive responses.

*Escherichia coli* encodes two NADH dehydrogenases, NDH-I and NDH-II, that together constitute the major entry point of electrons into the respiratory system. Simultaneous deletion of genes encoding these enzymes disconnects NADH oxidation from the respiratory chain, resulting in a severe growth defect and metabolic alterations. We performed adaptive laboratory evolution (ALE) of an NADH dehydrogenase-deficient strain to examine the compensatory redox network that enables *E. coli* to overcome the growth restrictions. We conducted a multi-omics analysis followed by genome-scale model-based calculations to delineate adaptive rewiring in the respiratory network that optimizes growth. We show that loss of NADH dehydrogenases initially forces cells into a fermentative metabolic state characterized by lactate secretion, likely to maintain PMF and redox balance. However, during growth optimization, the strains consistently select a bypass centered on malate:quinone oxidoreductase (Mqo), creating an alternative respiratory route that transfers electrons directly to the quinone pool. This bypass drives extensive reorganization of central carbon metabolism, restores respiratory electron flux, and promotes a pronounced shift from fermentative toward respiratory metabolism, thereby improving redox homeostasis and cellular fitness. Together, our findings reveal that bacterial respiratory systems have a remarkable metabolic buffering capacity to overcome failure of canonical respiratory pathways, necessitating a systems-level understanding for any potential therapeutic or translational targeting.

## Results and discussion

### Proximal impact of NADH dehydrogenase deficiency

To examine the immediate metabolic consequences of inhibiting NADH oxidation, we engineered an NADH dehydrogenase-deficient *E. coli* strain by knocking out the genes encoding NDH-I subunit B and NDH-II (Δ*nuoB*Δ*ndh*; hereafter ΔNDH). As expected, the lack of NADH dehydrogenase caused a pronounced growth defect, indicating that disrupting NADH oxidation imposes a substantial physiological challenge (Figure 1A) (6–8). Consistently, ΔNDH exhibited significantly increased lactate secretion, potentially due to D-lactate dehydrogenase (LdhA) mediated conversion of pyruvate to D-lactate, which regenerates NAD^⁺^ (Figure 1B) (8, 9, 19, 20). Notably, several fermenting bacteria use lactate secretion for metabolite ion-exchange-based PMF generation (21, 22). This compensatory response also reduces oxygen uptake rate (Figure 1C) (6). Thus, to mitigate the loss of NADH dehydrogenase activity, ΔNDH recruits a metabolically costly alternative strategy for NADH oxidation, reducing carbon utilization efficiency and energy conservation.

**Figure 1:**
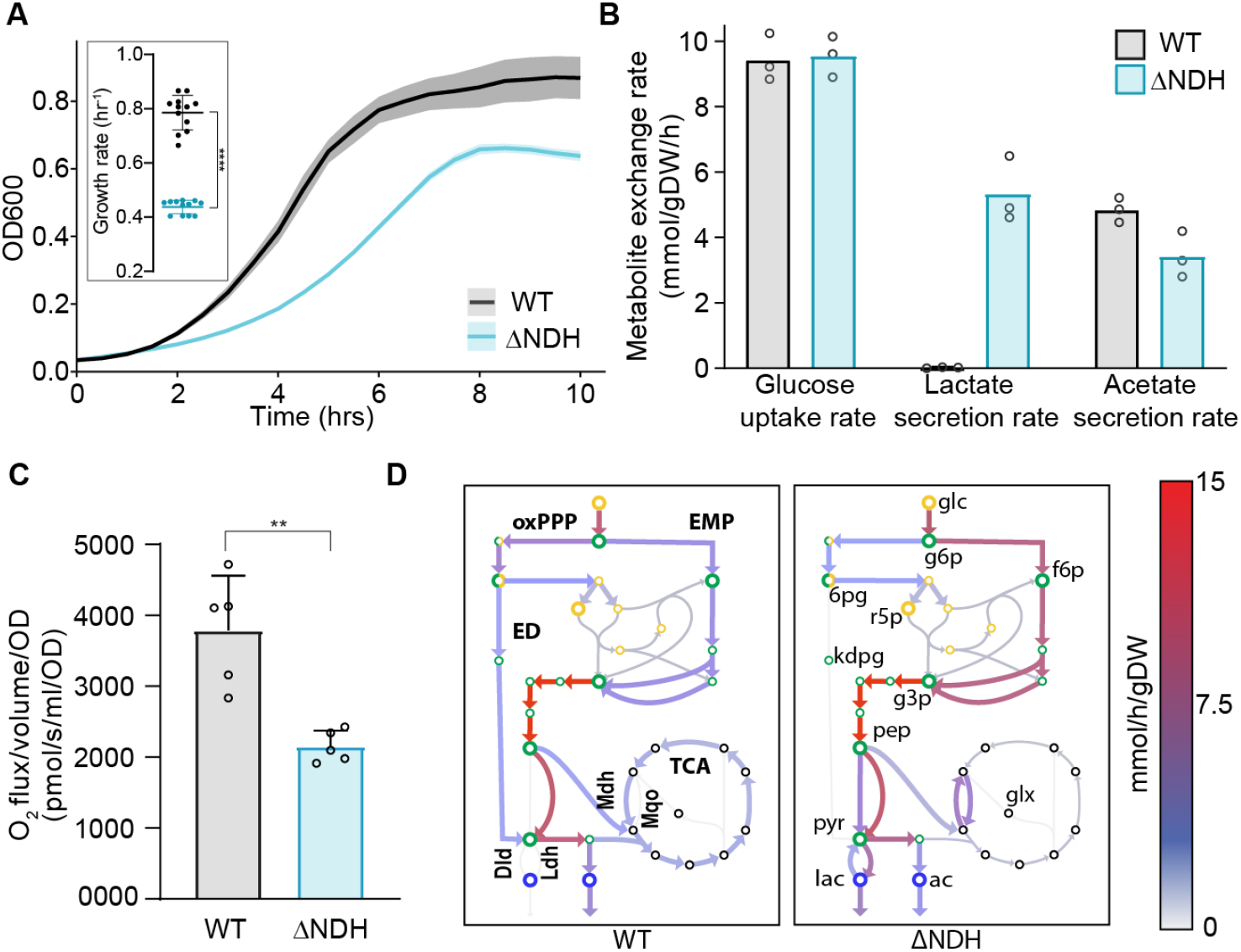
Proximal impact of NADH dehydrogenase deficiency. (A) Growth profile of wild-type (WT) and NADH dehydrogenase-deficient mutant (ΔNDH) of *E. coli* in M9 minimal medium supplemented with glucose under aerobic conditions. The growth curves show the mean of four biological replicates (with three technical replicates each). The shaded area shows the standard error of the mean. The inset shows the corresponding growth rate plot, where error bars represent the standard deviation. Statistical significance was calculated using an unpaired t-test (\*\*\*\**p*-value < 0.0001). (B) Metabolite exchange rate estimates of the WT and ΔNDH strains. Three biological replicates are depicted as individual points on the graph. mmol/gDW/h: millimoles per gram dry weight per hour. (C) Oxygen uptake rate measurements for WT and ΔNDH strains. Values for all five biological replicates are shown. Error bars represent standard deviation. pmol/s/ml/OD: picomoles per second per milliliter per optical density at 600nm. Statistical significance was calculated using an unpaired t-test (\*\**p*-value = 0.0019). (D) ME-model-based metabolic flux maps of the central carbon metabolism for WT and ΔNDH. Metabolites are indicated as follows: glc, glucose; g6p, D-glucose-6-phosphate; f6p, D-fructose-6-phosphate; 6pg, 6-phospho-D-gluconate; r5p, ribose-5-phosphate; kdpg, 2-keto-3-deoxy-6-phosphogluconate; g3p, glyceraldehyde-3-phosphate; pep, phosphoenolpyruvate; pyr, pyruvate; lac, lactate; ac, acetate; glx, glyoxylate. Enzymes: Dld, quinone-dependent D-lactate dehydrogenase; Ldh, D-lactate dehydrogenase; Mdh, malate dehydrogenase; Mqo, malate quinone oxidoreductase. Pathways: oxPPP, oxidative pentose phosphate pathway; EMP, Embden-Meyerhof-Parnas pathway; ED, Entner-Doudoroff pathway; TCA, tricarboxylic acid cycle. (Green empty circles: glycolysis nodes; Yellow empty circles: pentose phosphate pathway nodes; Black empty circles: TCA cycle nodes.)

The growth behavior and metabolite profiles of ΔNDH indicated large-scale metabolic rewiring. We therefore performed systems-level flux mapping with iJL1678b-ME, a genome-scale model of *E. coli* metabolism and protein expression (ME-model) (23). Each strain’s model was constrained with the measured glucose uptake, lactate and acetate secretion, and oxygen uptake rates. We layered the transcriptome as a constraint on protein allocation to pathways in central carbon metabolism (24, 25) (see Methods). Compared with the wild type, the ΔNDH strain showed a clear preference for the more energy-efficient glycolytic Embden–Meyerhof–Parnas (EMP) pathway (Figure 1D). This bias towards the EMP pathway over the Entner-Doudoroff (ED) pathway often results in higher ATP yield when aerobic oxidative phosphorylation is compromised (26, 27). Another striking observation was substantially reduced TCA flux in ΔNDH. Despite the decline in overall TCA cycle flux, we observed increased flux through both membrane-associated (Mqo) and cytosolic (Mdh) malate dehydrogenases in the ΔNDH strain (Figure 1D). Mdh catalyzes the thermodynamically favored NADH-dependent reduction of oxaloacetate to L-malate, whereas Mqo reoxidizes L-malate while reducing membrane quinones to quinols (20, 28, 29). Together, these enzymes enable a coupled system equivalent to an NADH dehydrogenase, oxidizing NADH and supplying electrons to the respiratory system. These metabolic adjustments likely help maintain NADH oxidation in the absence of the canonical respiratory route but do not restore optimal respiratory capacity, compromising the strain’s growth.

### Adaptive metabolic optimization for growth capacity restoration

Given NAD’s crucial metabolic role, we anticipated a layered capacity to maintain a growth-supportive NAD^+^/NADH balancing strategy. We performed ALE of the ΔNDH strain with four independently evolving lineages. All four lineages showed a significant improvement in growth rate in approximately 600 generations (Figure 2A). Such growth optimization is often based on selection of favorable genetic changes (30). Therefore, we performed whole-genome sequencing of the evolved ΔNDH strains and compared them with the genome sequence of the unevolved strain. In all evolved ΔNDH strains (eΔNDH A-D), we identified mutations that are commonly observed in the WT strain evolved to improve growth rate on glucose-supplemented minimal media (Supp. Table 1) (31). The eΔNDH-A and eΔNDH-D strains acquired mutations in RNA polymerase subunits that improve growth by enhancing RNA polymerase activity to promote overall transcription (31–33). In eΔNDH-B, eΔNDH-C, and eΔNDH-D, we observed intergenic mutations between the *pyrE* and *rph* genes that alleviate pseudo-auxotrophy in pyrimidine biosynthesis, leading to enhanced growth (31, 34).

**Figure 2:**
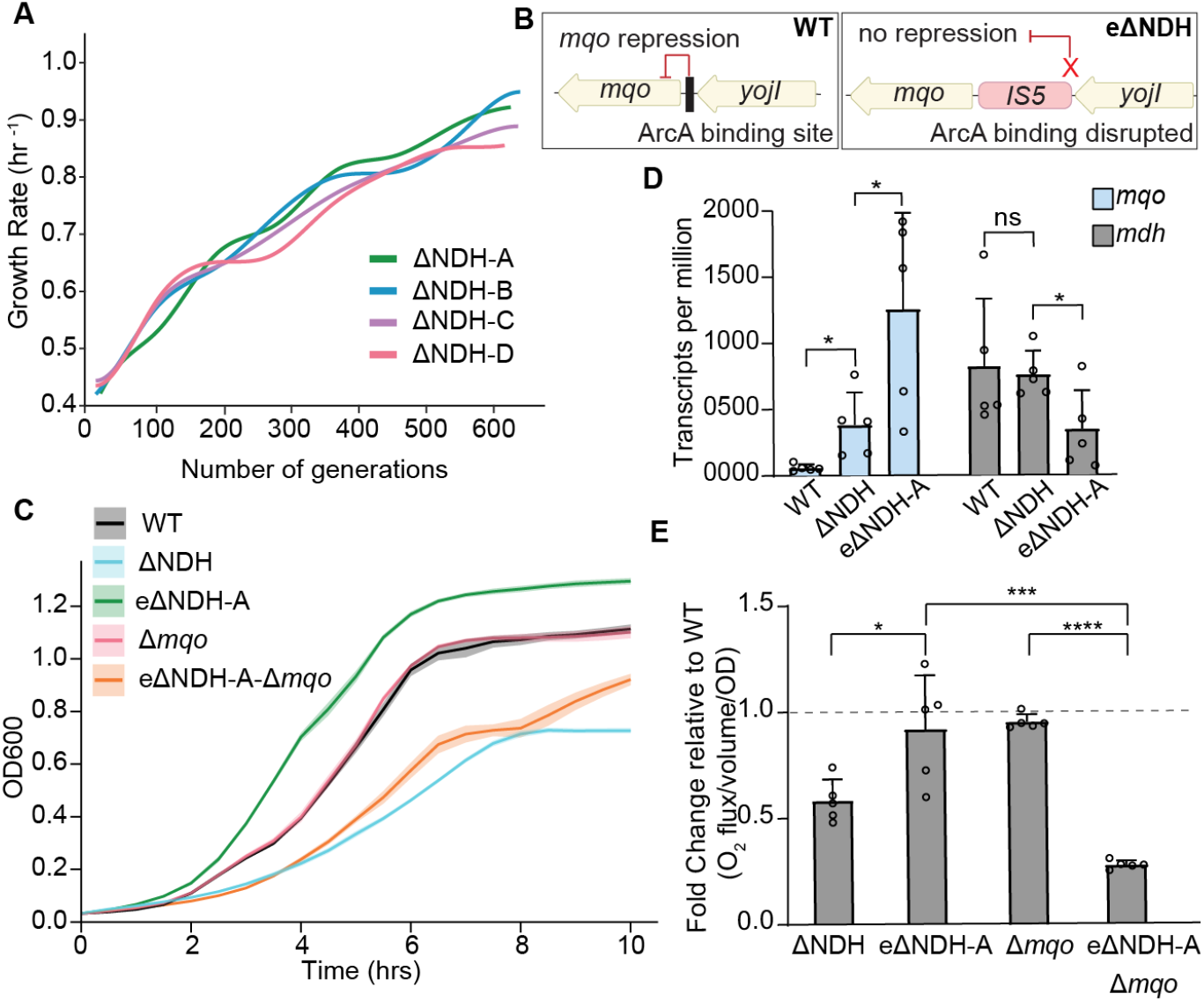
Adaptive laboratory evolution of NADH dehydrogenase-deficient strain. (A) Evolutionary trajectories depicting growth rate improvement of four independently evolving lineages of the NADH dehydrogenase-deficient mutant (ΔNDH A-D). (B) Schematic representation of the impact of the intergenic IS5 insertion on *mqo* expression regulation. The red cross in the eΔNDH scheme depicts the disruption of the ArcA binding site. (C) Growth profile showing the impact of *mqo* deletion on WT and eΔNDH-A. The growth curves show the mean of four biological replicates (with three technical replicates each). The shaded area represents the standard error of the mean. (D) Expression levels of *mqo* and *mdh* genes in WT, ΔNDH, and eΔNDH-A strains. Values for all five biological replicates are shown. Error bars represent standard deviation. Statistical significance was calculated using an unpaired t-test. (\**p*-value < 0.04) (E) Oxygen uptake rate of strains mentioned on the x-axis in comparison to WT. The dotted line shows the reference for WT level rate recovery. Values for all five biological replicates are shown. Statistical significance was calculated using an unpaired t-test. (\**p*-value < 0.0251, \*\*\**p-* value=0.0005, \*\*\*\**p*-value<0.0001)

Interestingly, all evolved ΔNDH strains acquired the IS5 insertion element upstream of the *mqo* gene (Supp. Table 1). The IS5 insertion occurs at the reported binding site of the transcriptional inhibitor ArcA, likely disrupting its binding and derepressing *mqo* (Figure 2B) (35). We therefore knocked out *mqo* in the eΔNDH-A strain to probe its adaptive involvement. While deleting *mqo* in the WT does not affect growth, knocking out *mqo* in the eΔNDH-A background causes a pronounced growth defect (Figure 2C). This validates the causal effect of the IS5 insertion upstream of *mqo* in optimizing eΔNDH-A growth. This gain-of-function nature of the *mqo* mutation was also supported by its significantly increased expression in eΔNDH-A compared with the unevolved ΔNDH (Figure 2D). While oxidizing malate to oxaloacetate, Mqo donates electrons to the ETS at the quinone level, thereby augmenting respiratory activity. We observed restoration of oxygen consumption in the eΔNDH-A strain, similar to the level of WT, which drops significantly when *mqo* is deleted from eΔNDH-A (Figure 2E). This increase in oxygen uptake rate was accompanied by a significant drop in lactate secretion in the evolved strain (Supp. Table 2). Thus, the ΔNDH strain’s growth limitation was a proximal response that was efficiently mitigated during adaptive evolution.

### Systems-level enantiostatic adaptation

Similar fitness can be achieved through distinct metabolic flux states available to a system. The fission yeast *Schizosaccharomyces japonicus* lacks oxic respiration but maintains efficient NADH oxidation, amino acid synthesis, and ATP generation by bifurcating the TCA cycle and optimizing glycolysis (36). In eΔNDH-A, EMP remained the preferred glycolytic pathway, as in ΔNDH. However, eΔNDH-A showed several notable metabolic distinctions (Figure 3A). The concerted function of quinone-dependent D-lactate dehydrogenase (Dld) with NAD-linked lactate dehydrogenase (LdhA) coupled NADH oxidation to respiration and prevented carbon loss as lactate (9, 20, 28, 37). We observed a significant replenishment of TCA flux in eΔNDH-A, likely because it partitioned the flux at the malate dehydrogenase node towards the canonical TCA cycle rather than restricting it to Mdh-Mqo cycling. Accordingly, we observed *mqo* upregulation in the evolved strain, but not a proportional change in *mdh* expression (Figure 2D).

**Figure 3:**
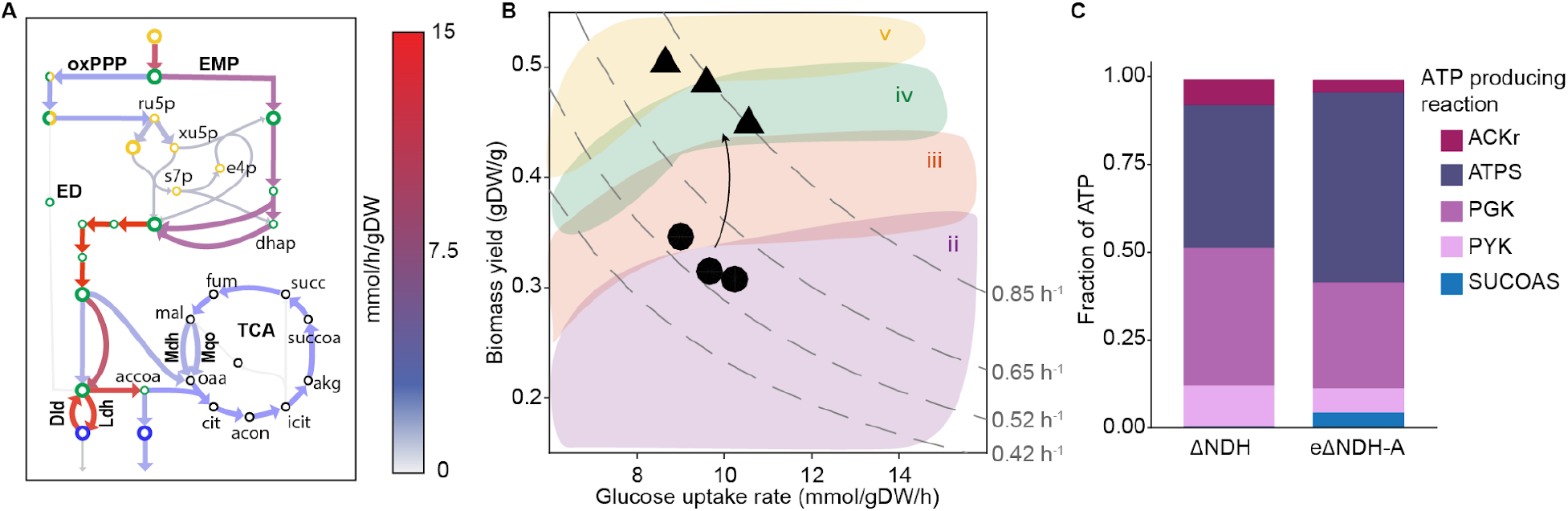
Adaptive re-routing of respiratory network. (A) ME-model-based metabolic flux map of the central carbon metabolism of the eΔNDH-A strain. Metabolites are indicated in the figure as follows: ru5p, D-ribulose-5-phosphate; xu5p, D-xylulose-5-phosphate, s7p, sedoheptulose-7-phosphate; e4p, D-erythrose-4-phosphate; dhap, dihydroxyacetone-phosphate; accoa, acetyl coenzyme-A; cit, citrate; acon, cis-aconitate; icit, icocitrate; akg, 2-oxoglutarate; succoa, succinyl coenzyme-A; succ, succinate; fum, fumerate; mal; malate; oaa, oxaloacetate. Enzymes: Dld, quinone-dependent D-lactate dehydrogenase; Ldh, D-lactate dehydrogenase; Mdh, malate dehydrogenase; Mqo, malate quinone oxidoreductase. Pathways: oxPPP, oxidative pentose phosphate pathway; EMP, Embden-Meyerhof-Parnas pathway; ED, Entner-Doudoroff pathway; TCA, tricarboxylic acid cycle. (Green empty circles: glycolysis nodes; Yellow empty circles: Pentose phosphate pathway nodes; Black empty circles: TCA cycle nodes) (B) Growth-yield plot showing aero-type classification of the ΔNDH (circles) and eΔNDH-A strains (triangles). Broken lines on the aero-type plot depict growth rate isoclines. Colored regions (ii, iii, iv and v) represent zones classifying different aero-types. Values of all 3 biological replicates are shown. (C) Relative contribution of ATP-producing reactions to total ATP generation in ΔNDH and eΔNDH-A strains. ATP production reactions depicted: (i) acetate kinase (ACKr) in mixed acid fermentation, (ii) ATP synthase (ATPS) in oxidative phosphorylation, (iii) phosphoglycerate kinase (PGK) and (iv) pyruvate kinase (PYK) in the glycolysis pathway, (v) succinyl-CoA synthetase (SUCOAS) in the TCA cycle.

The decrease in lactate secretion and increased oxygen uptake rate indicated higher reliance on aerobic metabolism in eΔNDH-A (Supp. Table 2; Figure 2E). The respiratory preference and associated cellular metabolic state influence the phenotypic positioning on the substrate uptake rate-biomass yield plane. This rate-yield coordinate provides a fitness descriptor, aero-type, representing the metabolic and phenotypic state, and the proteome complexity associated with a specific energy production scheme (38). Increased oxidative metabolism post-ALE shifted ΔNDH to a higher aero-type (Figure 3B). We observed a consistent shift in the source of ATP production. eΔNDH generated a higher proportion of its total ATP via ATP synthase than ΔNDH, with a fraction also coming from succinyl-CoA synthetase (Figure 3C).

### Concluding remarks

Combined deletion of both NADH dehydrogenases uncouples NADH oxidation from the respiratory chain, forcing immediate reliance on elevated Mdh-Mqo cycling to regenerate NAD^⁺^ alongside LdhA-mediated lactate fermentation. At physiological pH, lactic acid dissociates into lactate and hydrogen ions, which are symported to maintain pH homeostasis and consequently generate a proton gradient (39). Therefore, lactate production and its subsequent secretion in the unevolved ΔNDH strain likely offer dual metabolic benefits: (i) NADH oxidation via LdhA-mediated lactate production and (ii) PMF maintenance via lactate-proton symport. The proximal Mdh–Mqo cycling, however, is itself flux-limiting. The carbon shuttling between oxaloacetate and malate limits flux through the canonical TCA cycle, leaving the ΔNDH strain in a low-yield, respiration-restricted state. Adaptive evolution resolved this limitation by repartitioning flux at the malate node. Further upregulation of *mqo*, without a corresponding increase in *mdh*, shifted malate oxidation away from futile Mdh–Mqo cycling and toward the canonical TCA cycle, replenishing TCA flux while Mqo continued to feed electrons into the quinone pool. This repartitioning, together with reduced lactate efflux and a shift in ATP-generating flux toward oxidative phosphorylation, restored near-wild-type respiratory capacity. These findings motivate testing whether Mqo-associated bypasses operate in NADH dehydrogenase-dependent pathogens, where they could undermine single-target therapeutic strategies.

## Materials and method

### Estimation of NAD(P)H-associated reactions

The NAD(P)H-associated metabolic reactions were estimated from the iZY1689-StressME reaction set by identifying metabolic reactions containing NAD^⁺^, NADH, NADP^⁺^, or NADPH as explicit stoichiometric participants (40). Out of 2,296 unique metabolic reactions, 223 contained at least one of these cofactors (∼1 in 10).

### Bacterial strain generation and growth conditions

All experiments were performed with *Escherichia coli* K12 MG1655 (ATCC 700926) as the wild-type strain. P1 phage-based transduction method was used to generate knockout strains using Keio collection strains as donors for gene knockout cassettes (41, 42). Colony PCR and/or whole-genome sequencing were performed for strain verification (PCR verification primers are listed in Supplementary Table 3). Tecan Spark™ multimode microplate reader was used to perform growth characterization of all strains at 37°C with orbital shaking, using a 96-well plate with 200μL culture per well at atmospheric oxygen partial pressure. Each growth characterization experiment was performed in four biological replicates (each with three technical replicates). M9 minimal medium supplemented with 4 g/L glucose was used for planktonic bacterial growth. Growth rate calculations were done using gcplyr: an R package for microbial growth curve data analysis (43). The list of all reagents used in the study is provided in Supplementary Table 4.

### Adaptive laboratory evolution

Adaptive laboratory evolution (ALE) of the ΔNDH strain was performed with four independent lineages. Cultures were grown in M9 minimal medium supplemented with 4 g/L glucose at 37°C under aerobic conditions with continuous shaking. Cultures were serially propagated on fresh medium to maintain them in log phase. Cultures were always maintained under nutrient-rich conditions, as indicated by non-tapering exponential growth. The experiment was terminated once the strains reached a growth-rate plateau.

### DNA sequencing

Whole genome sequencing was performed for the end-point strains from each evolutionary lineage as described previously (44). Both clonal isolates and populations of every lineage were sequenced. Sequencing reads were filtered and trimmed using fastp software (45). Breseq version 0.38.1 was used to map sequencing reads and identify mutations relative to an *E. coli* K-12 MG1655 reference genome (NC_000913.3).

### RNA isolation and sequencing

For gene expression analysis, RNA was extracted from mid-log phase strains grown in M9 minimal medium supplemented with 4 g/L glucose at 37°C at 300 rpm. The harvested culture was pelleted down and dissolved in TRIzol (Thermo Fisher Scientific Cat. no. 15596018) and incubated at 65 °C for 15 min. RNA was obtained in the aqueous layer upon the addition of chloroform (SDFCL Cat. no. 3771OUR), which was then precipitated using chilled isopropanol (SDFCL Cat. no. 39694). This RNA pellet was washed with 70% and 100% ethanol (Honeywell Cat. no. 32221) and finally resuspended in Nuclease-free water (Invitrogen AM9932). The isolated RNA was further treated with DNase I (Invitrogen, Cat. no. AM2222) following the manufacturer’s protocol.

1 μg of RNA sample was used for rRNA depletion by NEBNext rRNA Depletion Kit (Bacteria) (Cat. no. E7850X), according to the manufacturer’s protocol. The NEBNext Ultra II Directional RNA Library prep kit for Illumina (Cat. no. E7760L) was used to construct double-stranded cDNA libraries, which were sequenced using an Illumina NextSeq 2000. Five biological replicates were sequenced for ΔNDH and eΔNDH-A. Three biological replicates were sequenced for WT. Two previously sequenced and available datasets for WT *E. coli* were batch-corrected and analyzed with the current three replicates (see data availability). RNA-seq data were analyzed as described previously (44), and the reads were mapped to the *E. coli* K-12 MG1655 reference genome (NC_000913.3). The batch correction of RNA-seq count data was done using ComBat-seq (46).

### Oxygen uptake rate measurement

The oxygen uptake rate estimation was performed by measuring the consumption of dissolved oxygen using the Oroboros O2K system. The cultures were grown in M9 minimal medium supplemented with 4 g/L glucose at 37°C and 300 rpm. Cells were harvested in their mid-log phase and diluted to a starting OD600 of 0.05. Oxygen uptake rates are either represented as oxygen flux per volume normalized to OD600 (pmol/ml/sec/OD) or as fold change in the oxygen uptake rate per OD600 relative to the WT.

### Metabolite exchange rate estimation

Strains were cultured in M9 minimal medium supplemented with 4 g/L glucose at 37°C under continuous shaking. Samples were collected at regular time intervals till stationary phase was reached. Culture density was measured at 600 nm absorbance with a spectrophotometer and correlated to cell biomass (dry weight). Cultures were filtered through a 0.22 μm filter (PVDF, Axiva, Cat no. SFPV13 R) for exometabolite estimation using High-Performance Liquid Chromatography (HPLC) (Agilent 12600 Infinity) with a Bio-Rad Aminex HPX87-H ion-exclusion column. The following method was used for HPLC: an injection volume of 10 μL and a 5 mM H_2_SO_4_ mobile phase set to a flow rate and temperature of 0.5 mL/min and 45°C, respectively, for a run time of 20 minutes per sample. A refractive index detector was used. Metabolite concentrations were determined using calibration standards prepared with different concentrations of a panel of metabolites dissolved in M9 minimal medium [Sodium-L-lactate (Sigma Aldrich Cat. No.71718), Sodium acetate (Sigma Aldrich Cat. No. S5636), and D-glucose (Sigma Aldrich Cat. No.49139), Sodium succinate di-basic (Sigma Aldrich Cat. No.14160), Sodium formate (Sigma Aldrich Cat. No. 798630) and Glycerol (MPI Cat. No. 1931996)].

### Metabolic flux distribution analysis

Flux mapping was done as previously described (24, 25) using iJL1678b-ME, a genome-scale model of *E. coli* K-12 MG1655 metabolism and protein expression (23). For ΔNDH and eΔNDH-A, both NADH dehydrogenases were removed by knocking out *ndh* (b1109) and *nuoB* (b2287) in the model. Each strain’s model was constrained with its measured glucose uptake, lactate secretion, and acetate secretion rates (means of three biological replicates). Expression data were layered on using the same method as the previous flux mapping (24). Transcript mass fractions of glycolysis, the oxidative and non-oxidative pentose phosphate pathway, and the TCA cycle were averaged over the five RNA-seq replicates of each strain and expressed as ratios to the ribosomal protein transcript mass fraction (47). Each ratio was imposed as a lower bound on the corresponding ratio of pathway protein to ribosomal protein in the model. Where the phenotypic data left the route of NADH oxidation undetermined, the protein allocated to the D-lactate dehydrogenases (LdhA and Dld) was also capped at a fixed multiple of its transcript-implied ratio.

Oxygen uptake was measured relative to WT, so the WT model was solved first with oxygen uptake unconstrained. The resulting WT oxygen uptake rate was then scaled by each knockout strain’s measured fold change (five biological replicates) and applied as a band of one standard deviation around the mean. At a given growth rate, the model returns the flux distribution that satisfies these constraints with the least total protein allocation. ME-models were solved with the quad-precision solver qMINOS through solveME (48, 49), and flux maps were drawn with Escher (50). Each reaction’s contribution to ATP production (Figure 3C) was calculated as its absolute flux divided by the summed absolute flux through the reactions of the aero-type definition (ACKr, ATPS4rpp, PGK, PPS, PYK, SUCOAS, PPK2r, PPKr and PRPPS) (38). The ATP synthase contribution is the aero-type.

### Quantification, visualization and statistical analysis

Quantification and generation of all plots were done using either Prism (GraphPad) version 11.1.0 or RStudio version 2024.12.0. All experiments in this study were performed at least in triplicate. All statistical tests were performed in Prism. The figure legends of each plot mention the details of the statistical tests used.

## Data availability

DNA sequencing and RNA sequencing data supporting this study are deposited in the NCBI SRA accession no. PRJNA1518820 and GEO accession no. GSE348946, respectively. The RNA-seq data for the additional two biological replicates of WT *E. coli*, WT-4 (GSM4036885) and WT-5 (GSM4036886), which were batch-corrected and analyzed with the current dataset, are available under GEO accession no.GSE135867.The strains generated in this project are available for distribution for research purposes.

## Code availability

Code for constraining and solving the ME-model, including the transcriptome-derived lower bounds and the upper bound on pathway protein allocation used here, is available at https://github.com/SBRG/ME-script (run_simulations_w_cap.ipynb). Simulations used COBRAme (23) and solveME (49).

## Author contributions

Conceptualization & supervision: A.A.; Funding: A.A.; Methodology: A.B., A.P., N.G., S.K., and A.A.; Investigation: A.B., A.P., and N.G.; Visualization: A.B., A.P., and A.A.; Writing the original draft: A.B. and A.A. All authors reviewed the manuscript.

## Acknowledgements

We thank Prof. Bernhard O. Palsson, University of California San Diego, for sharing resources for adaptive laboratory evolution. This work was funded by the Department of Atomic Energy, India-Tata Institute of Fundamental Research Grant (RTI4015) and Ramalingaswami Re-entry Fellowship, DBT, India (BT/RLF/Re-entry/70/2020) to AA.

## Supplementary Sheet 1

Metabolic reactions directly involving NAD(H) or NADP(H) in the *E. coli* StressME model.

## Supplementary information

Contains supplementary tables related to the study.

