## Supplementary Tables for "Adaptive respiratory bypass restores oxidative metabolism in NADH dehydrogenase-deficient *Escherichia coli*"

**Supplementary information for: Adaptive respiratory bypass restores oxidative metabolism in NADH dehydrogenase-deficient *Escherichia coli***

**Supplementary Table 1**. Mutations observed in evolved ΔNDH strains compared to the unevolved ΔNDH population. All mutations are listed with genomic locations in parentheses.

| Strains | Media specific mutations | Mutations common in all evolved lineages |
|---|---|---|
| eΔNDH-A | *rpoC* →<br>G→A<br>(4,185,642) | *mqo* ←/← *yojI*<br>IS5 (+) + 4bp<br>(+2,306,821) |
| eΔNDH-B | *pyrE* ←/← *rph*<br>Δ82<br>(3,815,859) | *mqo* ←/← *yojI*<br>IS5 (+) + 4bp<br>(+2,306,821) |
| eΔNDH-C | *pyrE* ←/← *rph*<br>Δ1<br>(3,815,810) | *mqo* ←/← *yojI*<br>IS5 (+) + 4bp<br>(+2,306,821) |
| eΔNDH-D | *pyrE* ←/← *rph*<br>Δ1<br>(3,815,810)<br>*rpoB* →<br>G→A<br>(4,183,258) | *mqo* ←/← *yojI*<br>IS5 (+) + 4bp<br>(+2,306,821) |

**Supplementary Table 2.** Metabolite exchange rates of ΔNDH and eΔNDH-A.

| | WT | ΔNDH | eΔNDH-A |
|---|---|---|---|
| Glucose uptake rate | 8.84, 9.15, 9.22 | 9.62, 10.14, 8.9 | 9.57, 10.7, 8.36 |
| Lactate secretion rate | 0.03, ND, ND | 4.9, 6.49, 4.61 | ND, 0.31, ND |
| Acetate secretion rate | 4.86, 5.26, 4.47 | 2.8, 3.29, 4.2 | 1.73, 3.4, 1.88 |

Values of all three biological replicates are depicted in the table. All units are millimoles per gram dry weight per hour. ND: Not detected.

**Supplementary Table 3.** List of primers

| Primer | Sequence (5'-3') |
|---|---|
| *Gene-specific primers* | |
| *ndh* FP | TGTTTTTTGATCTCACCCGG |
| *ndh* RP | TCCTTCTTTGCAGTTATGCC |
| *nuoB* FP | CATGGTCAACCTCTATCCGC |
| *nuoB* RP | TCCGGCCCAAAACGGTTGCG |
| *mqo* FP | GACTGCTGCCGTCAGGTCAA |
| *mqo* RP | TTGACAACGCGGTAGTTCGC |
| *mqo-IN-RP* | CGGATGCTTTACCGTAAACC |
| *Kanamycin cassette-specific primer* | |
| K1 (RP) | CAGTCATAGCCGAATAGCCT |

FP: Forward primer binding upstream of the gene; RP: Reverse primer binding downstream of the gene; IN-RP: Reverse primer binding internally in the gene.

**Supplementary Table 4: List of reagents**

| Reagent/resource | Source | Identifier or catalog number |
|---|---|---|
| $CaCl_2.2H_2O$ | Sigma-Aldrich | C3306 |
| $MgSO_4.7H_2O$ | SRL | 1344116 |
| $Na_2HPO_4$ | Sigma-Aldrich | S9763 |
| $KH_2PO_4$ | Sigma-Aldrich | P5655 |
| NaCl | MP Bio | 194848 |
| $NH_4Cl$ | Sigma-Aldrich | A4515 |
| $FeCl_3.6H_2O$ | Sigma-Aldrich | F2877 |
| $ZnCl_2$ anhydrous | Sigma-Aldrich | Z1052 |
| $CoCl_2.6H_2O$ | Sigma-Aldrich | C8661 |
| $Na_2MoO_4.2H_2O$ | Sigma-Aldrich | M1651 |
| $CuCl_2.2H_2O$ | Sigma-Aldrich | C3279 |
| $H_3BO_3$ | USB | 76324 |

| | | |
|---|---|---|
| Concentrated HCl | Sigma-Aldrich | H1758 |
| D-Glucose | Sigma-Aldrich | 49139 |
| Kanamycin | Sigma-Aldrich | K1377 |
| Luria–Bertani agar | Himedia | M1151 |
| Luria–Bertani broth | Himedia | M1245 |
| Sodium-L-lactate | Sigma Aldrich | 71718 |
| Sodium acetate | Sigma Aldrich | S5636 |
| Sodium succinate di-basic | Sigma Aldrich | 14160 |
| Sodium formate | Sigma Aldrich | 798630 |
| Glycerol | MPI | 1931996 |
| 0.22 µm PVDF filters | Axiva | SFPV13 R |
| TRIzol reagent | Thermo Fisher Scientific | 15596018 |
| Propan-2-ol Extrapure (isopropyl alcohol) | SDFCL | 39694 |
| Chloroform | SDFCL | 3771OUR |
| Absolute ethanol | Honeywell | 32221 |
| Nuclease-free water | Invitrogen | AM9932 |
| Nucleospin Tissue kit | Macherey-Nagel | 740952.50 |
| Nextera XT kit | Illumina | FC-131–1024 |
| DNase I | Invitrogen | AM2222 |
| NEBNext rRNA Depletion Kit (Bacteria) | New England Biolabs | E7850X |
| NEBNext Ultra II Directional RNA Library prep kit for Illumina | New England Biolabs | E7760L |
